# Optimizing methods for virome analysis based on studies of a synthetic viral community

**DOI:** 10.1101/2025.10.23.683462

**Authors:** Jiayi Duan, Andrew D. Marques, Matthew Hogenauer, Young Hwang, Yanjia Zhang, Aaron Timperman, Stephanie Higgins, Naomi G. Wilson, Elizabeth Aine Fitts, Haeun Karissa Lim, Kyle Bittinger, Ahmed M. Moustafa, Ronald G. Collman, Frederic D. Bushman

## Abstract

Studies of whole viral populations--the “virome”--are yielding exciting new insights into biological systems, but methods are still being optimized. Here we describe generation and use of a synthetic viral community to assess several technical challenges important in virome analysis. Our mock community was comprised of phages lambda, T4, M13, MS2, and phi6, together with adeno-associated virus (AAV), murine hepatitis virus (MHV), and vaccinia virus (VV). We spiked the mock community into different human sample types, including stool, saliva, oropharyngeal (OP) wash, and bronchoalveolar lavage (BAL), then passed the samples through different virus enrichment protocols and analyzed by Illumina sequencing. Compared to direct metagenomic sequencing, VLP enrichment protocols greatly increased viral read yields from virus-rich samples such as from stool and saliva. Three VLP enrichment work flows were compared, and each was found to have strengths and weaknesses. Four methods for DNA amplification were compared, with three showing over-amplification of small circular ssDNA viruses, most notably GenomiPhi. Studies of viral particle stability in the presence of nuclease showed that most viral genomes were stable when protected in viral particles, but phage MS2 RNA was unexpectedly labile under some of the conditions tested. Comparison of Illumina 1000-cycle sequencing versus 300-cycle sequencing showed that longer reads supported generation of longer viral genome assemblies. Bacteriophage DNA can be modified by at least 12 different chemistries, raising the question of whether these modifications might block recovery in virome analytical protocols. We tested bacteriophage T4 DNA modified with glucosyl-hydroxymethylcytosine (ghmC) and hydroxymethylcytosine (hmC), and found that both were readily detected, though the recovery of ghmC-modified DNA was reduced. These studies together with published data help provide guidance for virome researchers optimizing analytical protocols.

## Introduction

The planetary virome is vast. Virus-like particles (VLPs) on Earth are believed to number ∼10^31^[1-4]. Rich sea water can harbor ∼10^7^ VLPs per mL [5, 6]. Human stool can contain ∼10^9^ VLPs per gram [7-10]. Viruses can affect their hosts in numerous ways, both positively and negatively [9, 11-24]. Thus it is often of interest to characterize whole viral populations in a biome to assess the influence of these populations on function.

Because of the large numbers of viruses on Earth, an extremely small proportion have been studied in detail—thus in nucleic acid sequence data from a virome sample, many reads typically do not find close matches to anything in sequence databases. In addition, viral genomes typically evolve rapidly and so variants emerge quickly over time [25-27]. Thus, it is often difficult to know that sequences in “metagenomic dark matter” are viral in origin. Additionally, viral sequences often represent only a tiny fraction of nucleic acids in a biological sample, so the overwhelming majority of non-viral nucleic acids limit the ability to capture and identify viral sequences. Alternatively, when scant biological material including viruses is present, it may be challenging to capture viral sequences without enrichment.

For these reasons, it is often of interest to investigate the virome content in samples after enriching for VLPs before analysis by next-generation sequencing. In favorable cases, this will allow tentative identification of previously unstudied sequences as potentially virus-like in character. However, there are many different approaches to VLP enrichment, and many sample types of interest, so that no single method is ideal for all applications [8, 9, 11, 12, 18, 28-38].

Greatly complicating virome analysis is the heterogeneity of viral particles. Viral genomes can be comprised of RNA or DNA. Genomes can be single stranded or double stranded, linear or circular, and segmented or continuous. Viral particles can be of many morphologies, and enclosed in one or more lipid membranes (or not). Viral particles can range in size from 1.5 micrometers in length (pandoravirus) to 25 nanometers (Adeno-associated virus). Genome sizes range from 2.8 Mb for Pandoravirus to 2-3.9 Kb for Anelloviruses [4, 39, 40]. Genomes of satellite viruses and virus-like RNAs can be even smaller, comprised of structured circular molecules as little as 200-400 nt in length [27, 41, 42].

Here we describe steps toward optimizing methods for VLP enrichment based on assays with a synthetic viral community comprised of known viruses. Several previous publications have also investigated virome methods with spikeins of known viruses [18, 30, 32-35, 38, 43-46], which complement the work described here. As new virome research programs initiate, interest turns to capturing viruses that might have previously been lost during purification, motivating further rounds of method optimization. We included eight viruses in our mock community, named VirMock1, spanning a wide range of viral types (Table 1). Several questions pertinent to optimizing methods were addressed. Multiple methods for VLP enrichment have been reported (e. g. [8, 18, 28, 31, 33, 36, 37, 43-45])—how do these perform versus simple metagenomic DNA sequencing or RNA-seq on the same samples? When only small amounts of enriched viral genomes are available, amplification methods are often used— how much do these distort the relative abundances of viral genomes in a sample? How does the choice of sequencing method affect the data recovered? VLP fractions are often treated with nucleases to remove host cell nucleic acids—how frequently do these treatments also destroy viral genomes of interest? Bacteriophage DNA has been reported to be modified by at least 10 forms of covalent modification [47-49]—how often do these disrupt recovery in typical virome protocols? A complication is that many factors can influence the answers to these questions; here we report detailed surveys using our synthetic viral community to address some of the parameters influencing the output virome data produced. Optimized protocols are described in the methods section.

**Table 1.** Viruses included in the synthetic viral community VirMock1.

| **Virus** | **Enveloped?** | **RNA or DNA?** | **ss or ds?** | **Morphology** | **Genome size (bp)** | **Cell type supporting viral growth** |
| --- | --- | --- | --- | --- | --- | --- |
| T4 (wild-type with glyc-hmC) | no | DNA | ds | Tailed phage | 168,903 | *Escherichia coli* DH10B |
| lambda | no | DNA | ds | Tailed phage | 48,502 | *Escherichia coli* DH10B |
| pAAV-GFP | no | DNA | ss | icosahedral | 3,300 | Synthetic vector grown in human cell line |
| Vaccinia | yes | DNA | ds | brick shaped | 194,711 | BSC-1, african green monkey kidney cells |
| phi6 | yes | RNA | ds | spherical | 13,385 (all segments total) | *Psuedomonas syringae* LM2691 |
| MS2 | no | RNA | ss | icosahedral | 3,569 | *Escherichia coli* C3000 |
| MHV | yes | RNA | ss | spherical | 31,335 | 17Cl-1, murine fibroblast cell line |
| M13 | no | DNA | ss | filamentous | 6,407 | *Escherichia coli* C3000 |

## Results

### Comparison of methods for recovering VLP contigs

We initially compared viral sequence recovery after VLP enrichment versus simple metagenomic DNA or RNA sequencing of the unfractionated samples. For this, we first monitored just the viruses naturally occurring in the communities analyzed. Three methods were compared for VLP enrichment from the stool samples (Table 2), using methods derived from published protocols, here termed VP1 [44], VP2 [45] and VP3 [43]. All methods involved homogenization of stool in SM buffer, centrifugation to remove debris, filtration, nuclease treatment, and nucleic acid extraction. The methods differ in the details, including amount of starting material used, extent of initial dilution, filtration method, whether samples were concentrated after filtration, nature of the nuclease treatment, whether proteinase K treatment and phenol extraction were performed, and the nucleic acid purification kit chosen. Extracted RNA was subjected to cDNA sysnthesis and DNA and cDNA pooled and sequenced together. All three methods were tested on a common sample of homogenized stool (Fig. 1A). For comparison, we also analyzed viral sequence recovery from the same stool samples in pipelines based on conventional metagenomic DNA sequencing and total RNA-seq. Output sequencing reads were assembled into contigs and annotated for viral content using Sunbeam[50], then compared over the percentage of reads mapping to the viral contigs generated from the data.

**Figure 1.**
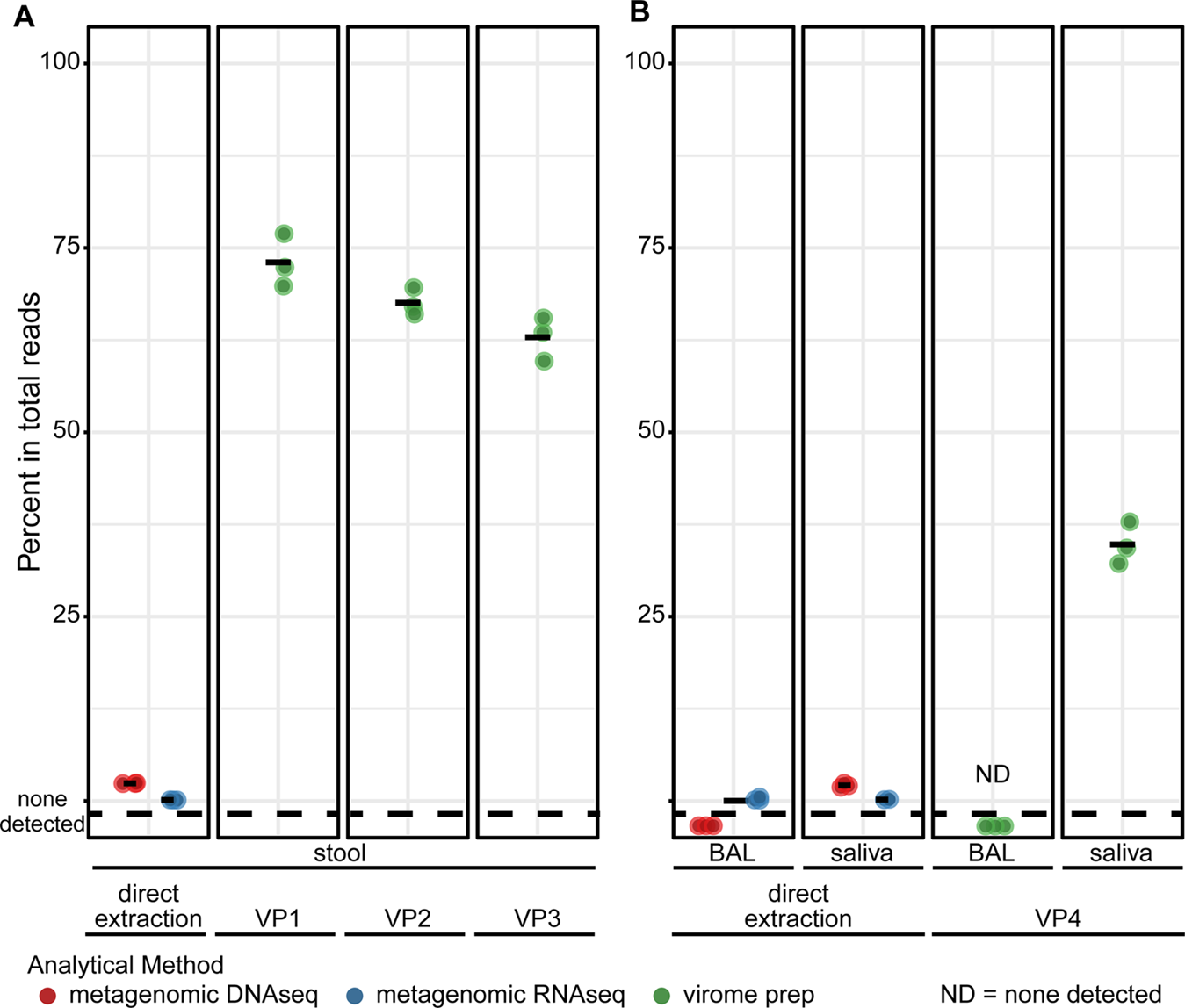
Comparison of viral sequence recovery after virus particle enrichment followed by sequencing, versus direct metagenomic sequencing of total DNA or RNA. A) Percentages of reads mapped to viruses annotated by Cenote-Taker2 in stool that was analyzed by direct extraction of DNA (red) or RNA (blue), or after viral particle enrichment and analysis using the VP1, VP2, or VP3 protocols (green). B) Percentages of reads mapped to viruses annotated by Cenote-Taker2 in saliva or BAL that was analyzed by direct extraction and metagenomic DNAseq (red) and RNAseq (blue), or by virus-like particle enrichment using the VP4 protocol (green). ND=no viral contigs detected.

**Table 2.**
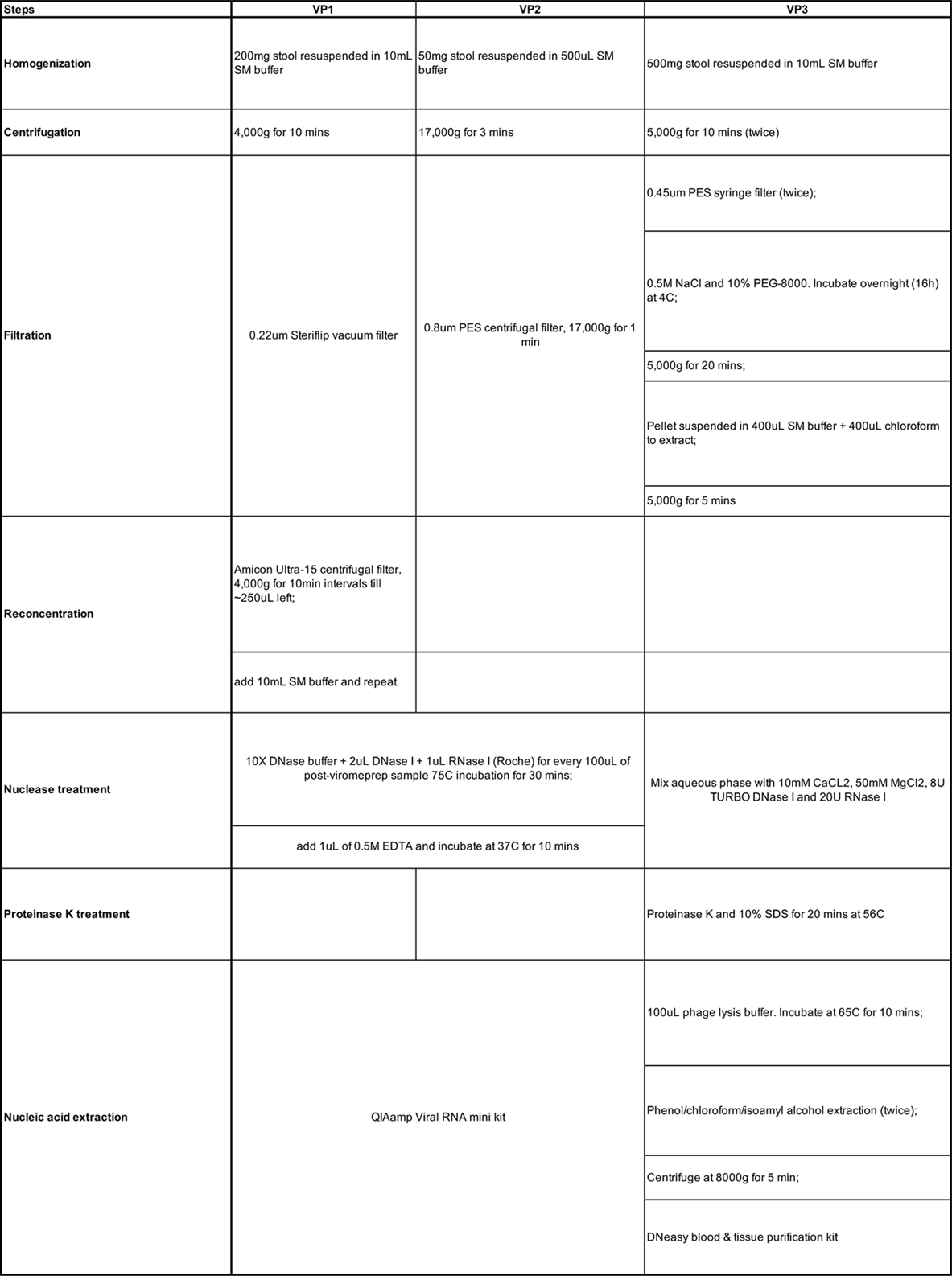
Steps in each of the three virus enrichment protocols.

Results with VP1-VP3 showed recovery of viral contig-aligning reads (DNA and cDNA combined) reaching an average of 62.9% to 73.0% of total reads, while metagenomic DNA- and RNA-seq returned only an average of 2.4% and 0.13%, respectively. The majority of recovered contigs annotated as Caudoviricetes, tailed dsDNA bacteriophages, in both virome sequencing and metagenomic sequencing (Fig. S1). Thus we conclude that VP1-VP3 are effective at enriching for virus-like particles and increasing recovery of virus-like contigs from stool.

We next carried out a similar comparison for saliva and BAL (Fig. 1B). The two represent samples types with lower biomasses than stool, with saliva still rich in viruses. BAL of healthy individuals, in contrast, is quite sparse for viruses, cells and biological materials generally. For analysis of these liquid samples, an enrichment protocol was devised paralleling VP1 but with reduced sample dilution and use of filters with larger pore sizes, which was added in an attempt to accommodate the higher viscosity of some airway samples (named VP4.

Saliva undergoing VP4 had on average 34.8% of sequencing reads annotated as viral. Saliva analyzed using metagenomic DNAseq and RNAseq without enrichment yielded only 2.1% and 0.179% viral reads on average, respectively. For BAL, VP4 resulted in no recovery of viral contigs, possibly due to losses from sample dilution and handling, and the sparse viral populations in the starting material. Processing of samples with quite low biomass through the VLP enrichment protocols, including BAL, oropharyngeal swab eluates and nasopharyngeal swab eluates, commonly resulted in sample loss (data not shown), indicating that other analytical approaches may be needed for some low biomass samples.

To ask how these methods perform with viruses of widely differing particle compositions, we next turned to tests of recovery of known viruses spiked in to human-derived samples.

### Assembling a synthetic viral community

Viral particles have an extremely wide range of genome sizes, and genomes can be comprised of RNA or DNA, single- or double-stranded, linear or circular, and segmented or continuous. Contemporary virome studies seek to capture all of these virus types while minimizing host and environmental contamination. As a tool for methods optimization, we thus assembled a community with a wide range of virus types, where each virus was well characterized and straightforward to grow.

Viruses used in our synthetic community, named VirMock1, are summarized in Table 1. We included viruses with RNA genomes (phi6, MS2, and MHV) and DNA genomes (AAV, lambda, M13, T4, and VV), with both single-stranded and double-stranded representatives. Particles had multiple morphologies, including enveloped and nonenveloped. Viral mixtures were characterized for infectivity and for numbers of viral genomes per mL using pPCR.

To analyze an example of unusual covalent DNA modification, we included phage T4 and two mutant derivatives (Table S1 and see below). In wild-type T4, about half of all cytosines are reported to be modified to glycosylhydroxymethylcytosine, which can block attack by host cell nucleases [49, 51, 52]. T4 mutants are available which only modify genomic DNA to hydroxymethylcytosine, or which incorporate cytosine only. Effects of these forms of DNA modification on virome recovery and sequencing protocols are assessed below. In our standard VirMock1 cocktail, we used wild-type T4 only.

To test virome recovery using VP1-VP3, we pooled these viruses together and spiked the VirMock1 community into a common stool sample or into SM buffer. Purified virome nucleic acids were analyzed using Illumina short read sequencing. Sequence reads were 1) directly mapped to genomes of VirMock1 community members to quantify VirMock1 recovery, and 2) used to generate contigs, followed by contig annotation, and then mapping of reads back onto viral contigs to quantify all viruses in the samples. Each sample type was analyzed in triplicate. This allows assessment of the results of different analytical approaches in a context where we have a clear expected answer.

Figure 2 presents results for viral detection and proportions of viral sequence reads recovered. Figure 2A quantifies the relative proportions of reads of VirMock1 and total viruses annotated; Figure 2B shows the relative proportions of the VirMock1 community members recovered; Figure 2C shows annotation of all viruses detected in the stool samples.

**Figure 2.**
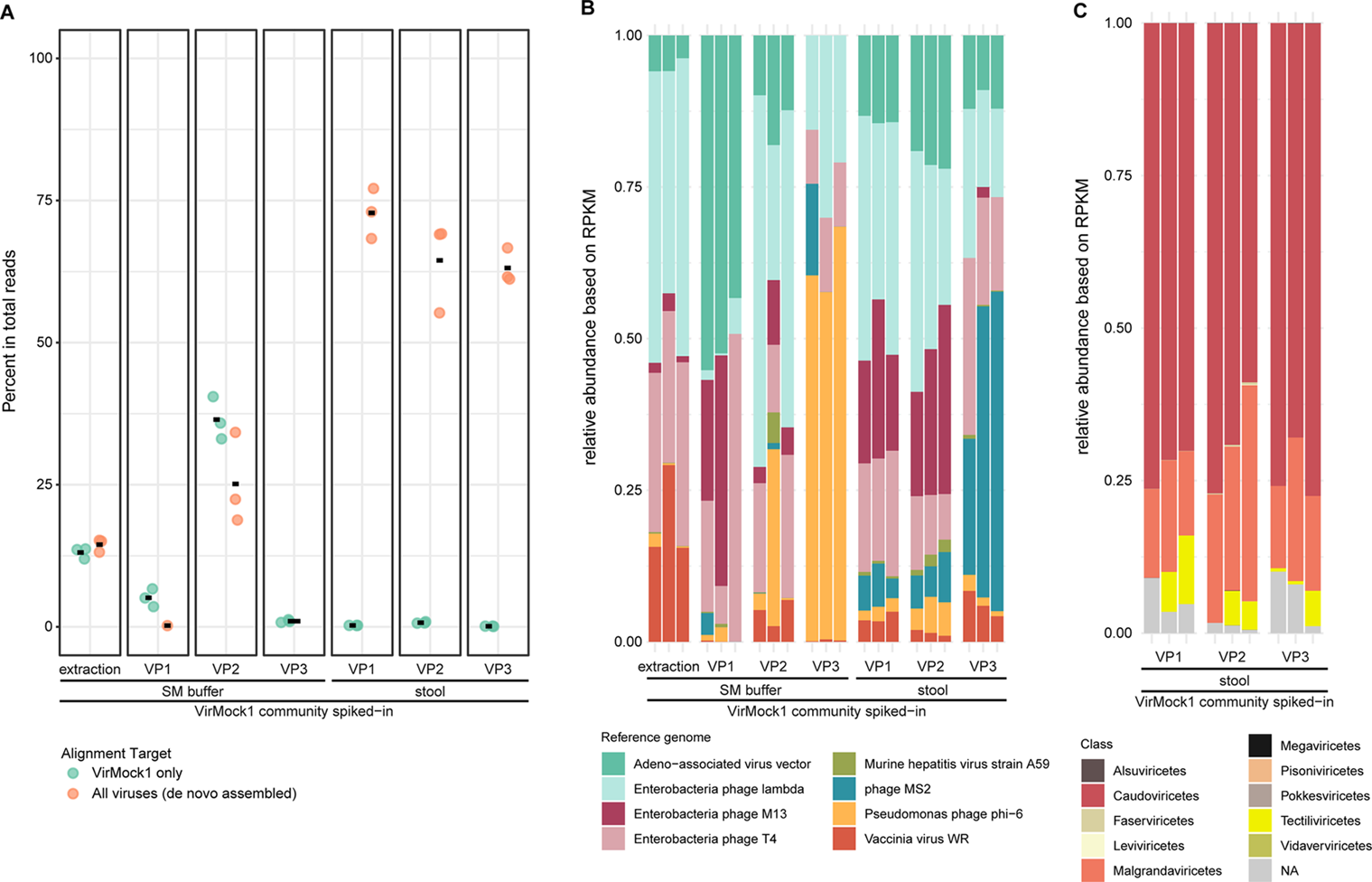
Comparison of methods for purifying viral particles from stool. A) Relative abundance of each reference virus in samples where VirMock1 was spiked into SM buffer or stool. Samples were directly extracted for nucleic acid, or first enriched for viral particles using VP1, VP2, or VP3. “Extraction” indicates samples directly extracted without any preceding particle enrichment. Reads aligning to any virus as annotated by Cenote-Taker2 are indicated in tan, reads annotating as aligning to a virus in VirMock1 are shown in green. B) Percentage of reads mapping to viruses from VirMock1 after VirMock1 was spiked into SM buffer or stool. Samples were directly extracted for nucleic acid, or first enriched for viral particles using VP1, VP2, or VP3. C) Relative abudance of classes of viruses as annotated by Cenote-Taker2 in stool after enrichment for for viral particles using VP1, VP2, or VP3.

As a reference, the VirMock1 community was mixed with SM buffer, treated with nuclease, and directly extracted (Fig 2A and B; left set of bars). After direct extraction, 13.1% of reads aligned to the VirMock1 genome sequences (Fig. 2B, left side, blue dots). Assembly of all viral contigs and mapping of reads back to contigs yielded a similar result (tan dots).

A similar analysis was performed after carrying out the VP1-VP3 purification protocols on VirMock1 spiked into a single common stool sample (Fig. 2A, right 3 panels), or spiked into SM buffer (Fig. 2A, panels 2-4). After spiking into the stool sample, for VP1, an average of 72.8 (± 4.41)% of reads mapped to viral contigs; numbers were slightly lower for VP2 (64.5 ± 8.00%) and VP3 (63.1 ± 3.06%). The proportions of the VirMock1 sequences in these samples were much lower, in the range of 0.1% to 0.8% of all reads, but this reflects dilution by the large numbers of viral sequences authentically present in the stool samples and recovered after viral enrichment.

In contrast, the recovery of VirMock1 sequences was more variable after spiking in to SM buffer only. VP2, the method involving the least dilution of the sample, showed the best recovery, while heavy losses of VirMock1 were seen with VP1 and VP3. This comparison further indicates that dilution can impair recovery, and that biological materials such as stool can improve recovery, potentially via “carrier” effects.

Figure 2B shows the VirMock1 lineages detected in each sample as read out from the Illumina sequencing data. Direct extraction of VirMock1 treated only with nuclease (DNAse I and bovine pancreatic RNase) before purification showed robust representation of six of the VirMock1 viruses, but two RNA viruses, phage MS2 and murine hepatitis virus, were present in only trace amounts. All eight VirMock1 viruses were detected after spiking into stool, again suggesting a possible “carrier” effect of the biological sample, in which the presence of stool components may stabilize labile viruses and facilitate their recovery.

Figure 2C shows the annotation of all viral contigs in each stool specimen at the Class level. The samples of stool with VirMock1 spiked in were dominated by Caudoviricetes, an abundant lineage of tailed bacteriophages with double stranded DNA genomes, and Malgrandaviricetes, a group containing common single stranded DNA bacteriophages. Both of these Class are commonly reported from analyses of bacteriophages in human stool [13, 24, 36, 53-59].

### Distortions of recovered communities associated with different DNA amplification methods

DNA amplification is frequently employed to allow analysis of samples with low microbial biomass. Therefore we next tested how different DNA amplification methods affected virome recovery and inferred compositions of the VirMock1 defined community. We assessed three types of VirMock1 preparations for this experiment: VirMock1 1) spiked into saliva and purified via VP4, 2) spiked into SM buffer to a final volume of 600 μL and purified via VP4, or 3) spiked into SM buffer to a final volume of 600uL, nuclease-treated, and directly extracted. Samples were then reverse-transcribed and amplified with one of four commercially available amplification kits: GenomiPhiV3[60], Primary Template-directed Amplification (PTA)[61], Whole Transcriptome Amplification (WTA2)[62], or Mulptiple Annealing and Looping-Based Amplification Cycles (MALBAC)[63] (Table S2) prior to Illumina DNA sequencing. MDA is an exponential amplification method; PTA, WTA2, and MALBAC are quasi-linear. We also included analysis of VirMock1 with no amplification. Samples were analyzed in triplicate to allow an evaluation of variance.

In the VirMock1-spiked SM buffer samples treated with nuclease and directly extracted without amplification (Fig. 3A, top right), the sequencing result generally reflected the input VirMock1 composition (Table S3). Samples showed good recovery of six of the VirMock1 viruses, and trace amount of phi6 and MS2. Roughly similar patterns were seen for the VP4-purified unamplified samples (Fig. 3A, top, left and middle), except that MHV and AAV showed increased representation after purification.

**Figure 3.**
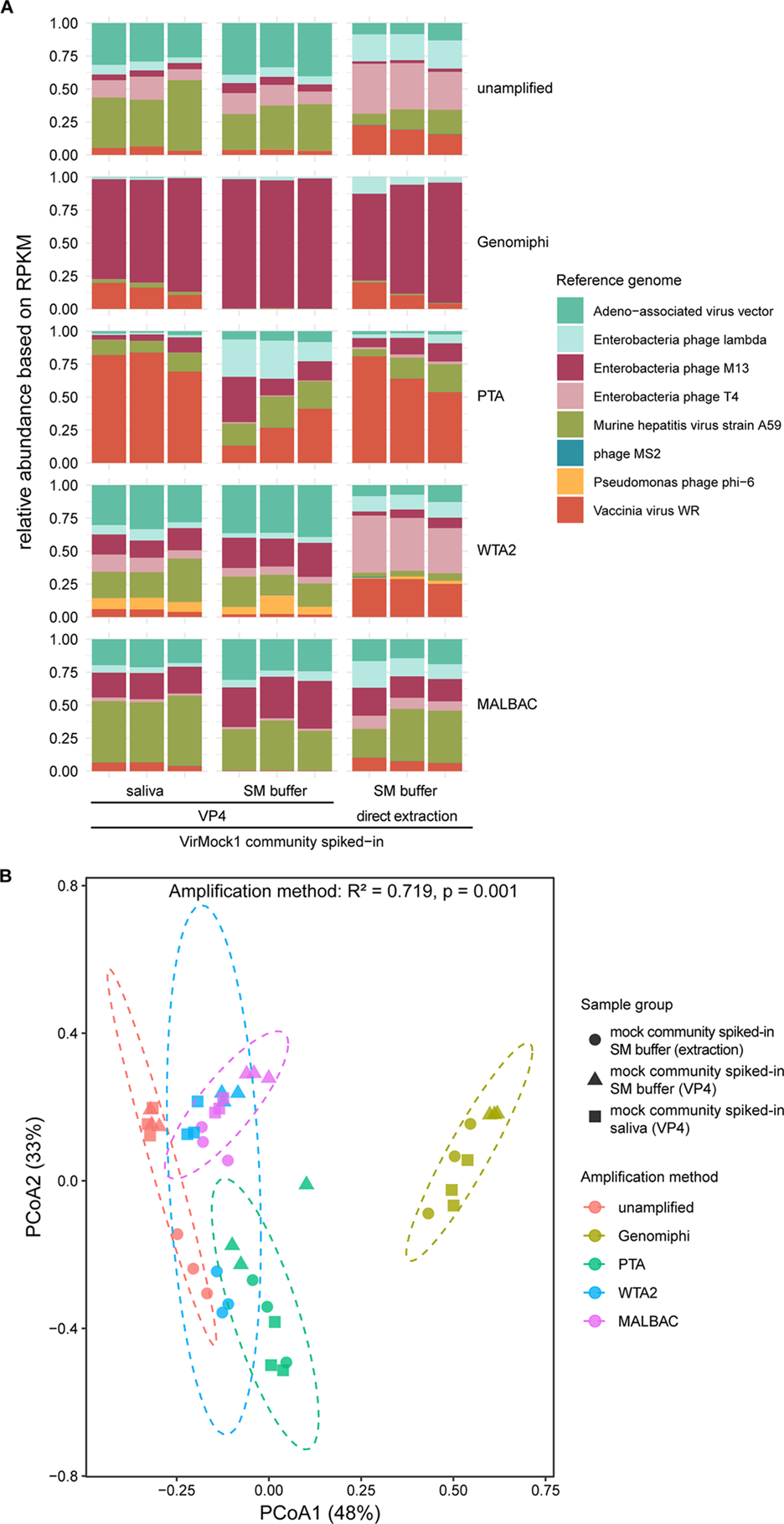
Comparison of the effects of different DNA amplification methods on viral genome recovery. A) Relative abudance of each VirMock1 reference virus; samples were compared after addition of VirMock1 to saliva or SM buffer and purified using a modified protocol with elements of VP1 and VP2 optimized for saliva (VP4). Samples were also tested after spiking into SM buffer and purification immediately afterwards. Samples were analyzed by Illumina sequencing without amplification, and after amplification using GenomiPhi, PTA, WTA2 with 17 PCR cycles, or MALBAC with 17 PCR cycles. B) PCoA analysis was performed using Bray-Curtis distances computed based on relative abundance of VirMock1 reference virus in each sample. Variation in community composition across amplification methods was assessed using PERMANOVA. Amplification methods (represented by different colors) and samples (represented by different shapes) were as in A.

Different amplification methods resulted in different degrees of enrichment of viral genomes and showed different biases for the various VirMock1 community members. GenomiPhi greatly favored M13 (Fig. 3A, second row) probably reflecting rolling circle replication of the small circular ssDNA genome, as has been reported in several previous studies[32, 64, 65]. The other three methods showed less extreme distortions of the viral populations (Fig. 3A, 3^rd^-5^th^ rows). PTA did not show such overamplification of M13, but resulted in considerable amplification of vaccina virus (Fig 3A, third row). The reason for favored amplification of vaccinia DNA is unknown. PTA, WTA2, and MALBAC all showed lower and variable levels of overampification of M13 in some conditions (Fig 3A and Fig. S2). Amplification using MALBAC showed overall similar relative abundance of VirMock1 viruses compared to unamplified, except for a moderate bias favoring MHV and M13 (Fig. 3A).

Figure 3B summarizes these data as a principal coordinate analysis based on Bray-Curtis distances between the measured compositions of samples. As can be seen, samples tended to cluster by amplification method. Genomiphi was the most extreme outlier, with the other methods closer to unamplified but clustering separately. Thus we conclude that the GenomiPhi method is too biased for routine use, and the other methods recovered most viruses effectively, though each has it’s own modest biases.

### Virus types differ in their sensitivity to nuclease treatments

VLP enrichment methods typically involve a nuclease step following VLP enrichment and before nucleic acid purification, intended to help remove free nucleic acids[8, 16, 18, 19, 28, 30-33, 35-38, 46, 53, 54, 64, 66-68]. However, such a step has the danger of degrading nucleic acid within particles if there is any degree of accessibility of encapsulated viral genomes[8, 66, 67, 69]. We thus incubated VirMock1 in SM buffer with a titrated mixture of DNAse and RNase, and quantified viral genome recovery after extraction. Quantification was carried out using Illumina DNA sequencing (Fig. 4A) or qPCR (Fig. 4B). We also compared results after spiking VirMock1 into stool (Fig. 4C). Recovery of each virus in VirMock1 was quantified relative to the measured input.

**Figure 4.**
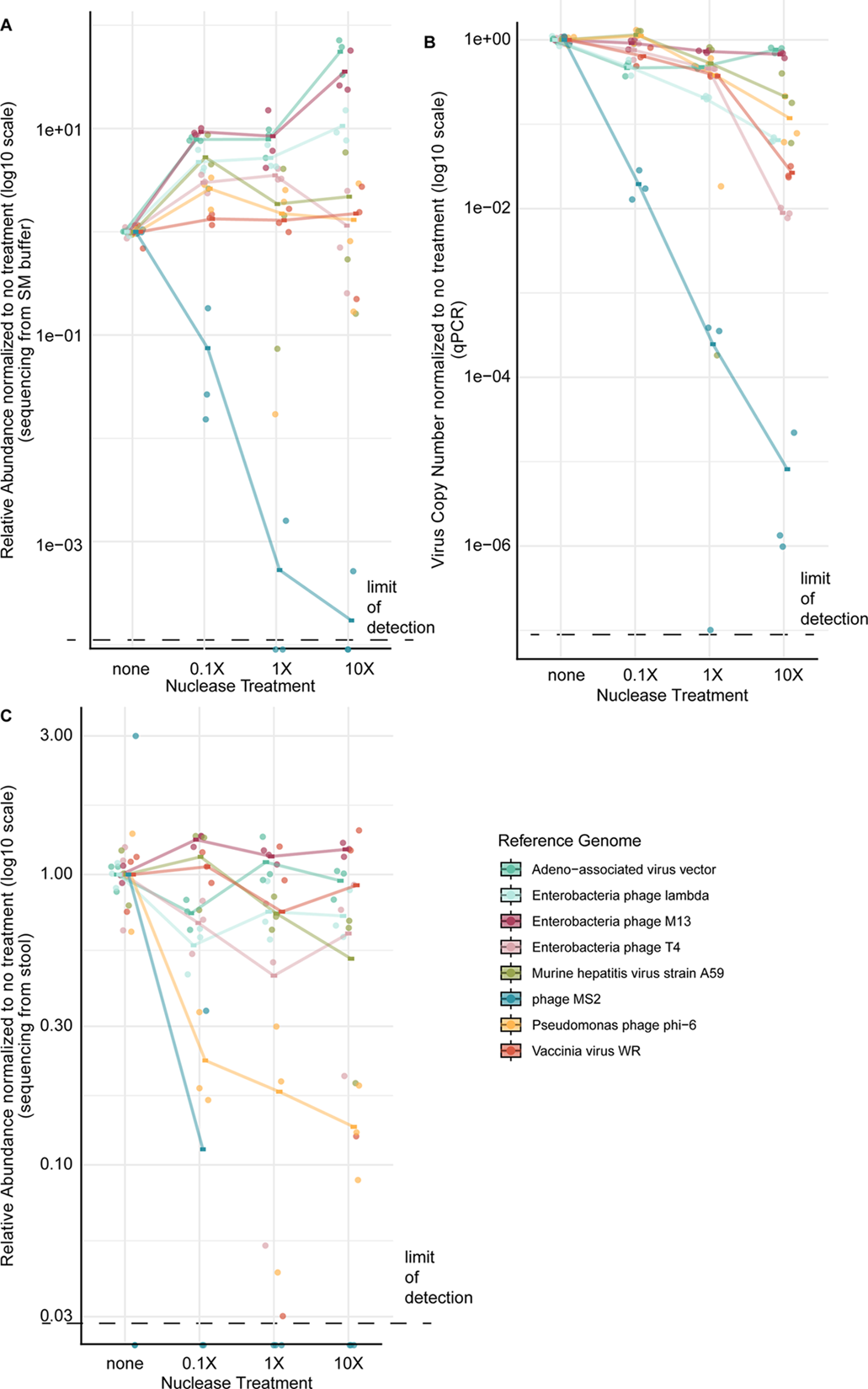
Assessing the stability of viral genomes within viral particles in the presence of nucleases. Nuclease treatment titrated from no added nuclease to 10X the protocol value was performed in the presence of VirMock1. Both RNase and DNase were added. In each case, relative abundances were normalized to the mean of the no-treatment control triplicates. Trendlines are plotted using the mean of triplicate measurements. The dashed line indicates the limit of detection in each assay. A) Relative abundance of each virus after nuclease treatment calculated based on their respective RPKM determined after Illumina sequencing. B) Nuclease titration results determined from qPCR for each VirMock1 viral genome. C) VirMock1 was spiked into stool, different amounts of nuclease added, and genomes recovered and quantified using RPKM after Illumina sequencing.

Recovery of viruses varied among types. Most viruses were insensitive to nuclease up to 10X the concentration used in typical VLP preparations. An exception was bacteriophage MS2, which was sharply reduced in abundance by both measures at even 0.1X nuclease concentration. MS2 was also lost when spiked into stool in two out of three replicates even without nuclease treatment, perhaps due to handling or pre-existing nuclease activity in stool (Fig 4C). Other viruses showed less susceptibility to nuclease; phi6 showed some decrease in relative abundance when spiked into stool (Fig 4C). Control experiments showed that free nucleic acids were reduced in abundance efficiently by 1X nuclease treatment when passed through VP4 (Table S4). Thus we find that most viruses withstood the nuclease treatment robustly, but one non-enveloped linear RNA virus (MS2) was quite sensitive.

### Comparing Illumina short read versus longer read sequencing

To begin to examine how the choice of sequencing method affects the output contig length, we compared the new Illumina 1000-cycle kit and the standard 300-cycle kit. For this, we used the same sequencing libraries generated from a common stool sample spiked with VirMock1 and purified using VP1 (Fig. 5). Samples were sequenced and analyzed using the contig assembly and annotation pipeline described in the methods section.

**Figure 5.**
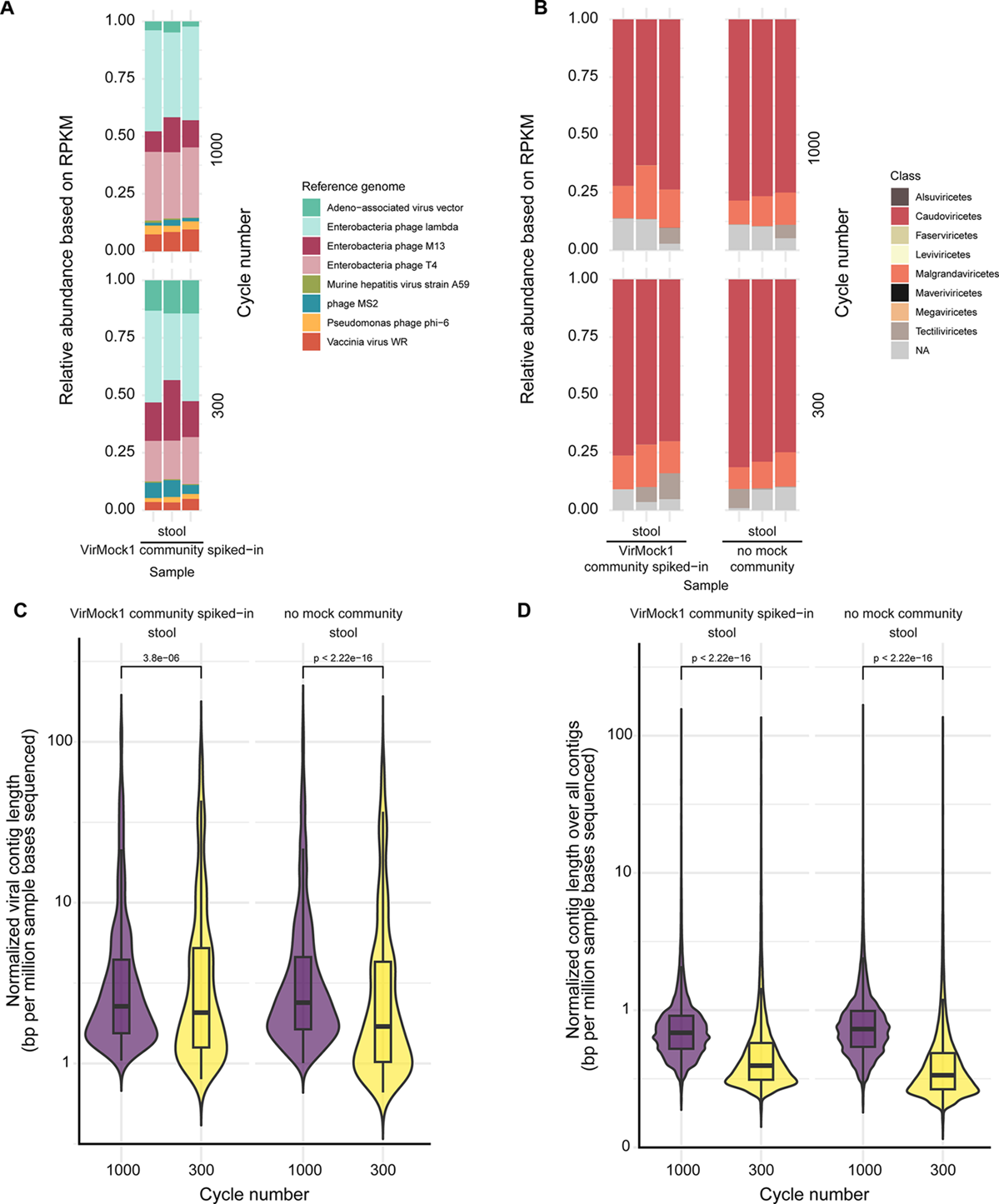
Comparison of Illumina sequencing with the Illumina 1000-cycle kit and 300-cycle kit platforms. A) Relative abundance of each reference virus in VirMock1 spiked-in stool that underwent VP1 and was then sequenced using the 1000-cycle kit (upper set of bars) or the 300-cycle kit (lower set of bars). B) Relative abudance of classes of viruses as annotated by Cenote-Taker2 in stool with or without VirMock1 spiked-in that underwent VP1 and was then sequenced using the 1000-cycle kit (upper set of bars) or the 300-cycle kit (lower set of bars). C) Wilcoxon rank-sum test was performed to compare the density of the length of the viral contigs output by the 1000-cycle kit and 300-cycle kit. D) Wilcoxon rank-sum test was performed to compare the density of the length of all contigs (not only viral) output by the 1000-cycle kit and 300-cycle kit.

The proportions of virus types in the stool samples were similar between the 1000-cycle kit and 300-cycle kits for both the VirMock1-spiked and unspiked samples (Fig. 5 A and B). Mock community members were detected after spike in of VirMock1, and an unbiased virus annotation emphasized the presence of Classes Caudoviricetes and Malgrandaviricetes as expected for stool samples.

We next compared the distribution of contig lengths returned after use of each sequencing method to assess whether the 1000-cycle kit generated longer contigs. For comparison, we normalized the length of each contig by the total number of bases sequenced in that sample. Given the same number of bases sequenced, the 1000-cycle kit resulted in more total contigs and viral contigs of longer length than the 300-cycle kit. This was true for both the VirMock1-spiked and unspiked stool samples (Fig. 5 C and D). The median length of viral contigs assembled from spiked and unspiked stool was 576 and 587 bp (2.27 and 2.39 bp per million sample bases sequenced) using the 1000-cycle kit, and 449 and 450bp using the 300-cycle kit (2.07 and 1.70 bp per million sample bases sequenced), respectively (Table S5). Thus, use of the longer length Illumina sequencing platform did result in longer length contigs (p < 0.001).

### Beginning to assess the possible influence of covalent DNA modification

Viral nucleic acids are subject to numerous covalent modifications, which have the potential to disrupt recovery of viral genomes in metagenomic analysis. Bacteriophage DNA has been reported to be subject to at least ten forms of DNA modification [47-49]. RNA broadly has been reported to be subject to >100 forms of covalent modification [70].

As one step toward assessing how much these modifications may disrupt metagenomic analysis of viral genomes, we compared three genotypes of bacteriophage T4: wildtype T4 (termed T4ghmC), T4hmC, and T4C. For wild-type T4, >50% of C residues are modified to glucosyl-hydroxymethylcytosine (see below). T4hmC is a mutant strain that produces only hydroxymethylcytosine. T4C harbors mutations in genes required for substituting HMC for dCTP and thus only contains unmodified cytosines in genomes. The effects of these modifications were previously documented in DNA sequencing studies as effects on inter-pulse distances in Pacific Biosciences DNA sequencing [49].

To test the effects of these modifications in our Illumina sequencing protocol, we first verified the presence of each DNA modification in the three T4 strains. We used LC-MS/MS to verify that the expected derivative of C predominated for each strain (Fig. S3). We also purified each DNA and showed that sensitivity to restriction enzyme digestion was as expected—that is, DNA containing glucosylhydroxymethylcytosine and hydroxymethylcytosine were protected from digestion by Alu I, while DNA containing cytosine was sensitive (Fig. S4). These studies documented the predominance of the expected forms of cytosine in each T4 strain.

We then asked whether these modifications disrupted genome sequencing. We mixed each form of T4 DNA with bacteriophage lambda DNA, which lacks the above modifications, and compared recovery after extraction, library preparation and DNA sequencing (Fig. 6). When mixed with lambda, the relative abundance of T4ghmC assayed by sequencing was more than 3-fold lower than that expected from the qPCR values for total genomes. For T4C and T4hmC, when mixed with lambda, there was no major reduction in relative abundance by sequencing compared to quantification by qPCR (Fig 6A, Table S6). Thus, hydroxymethylation does not detectably inhibit detection by sequencing, but glucosyl-hydroxymethylation did appear to diminish detection modestly. Similar conclusions were reached by Zhai et al.[38] Given that the Nextera method was used to prepare the sequencing library, we conjecture that the transposon-based step used to introduce amplification primers may be partially inhibited by glucosylhydroxymethylation, though the modified genomes were nevertheless detectable in our samples. It will be useful to document effects (if any) of further forms of DNA and RNA modification on virome assays.

**Figure 6.**
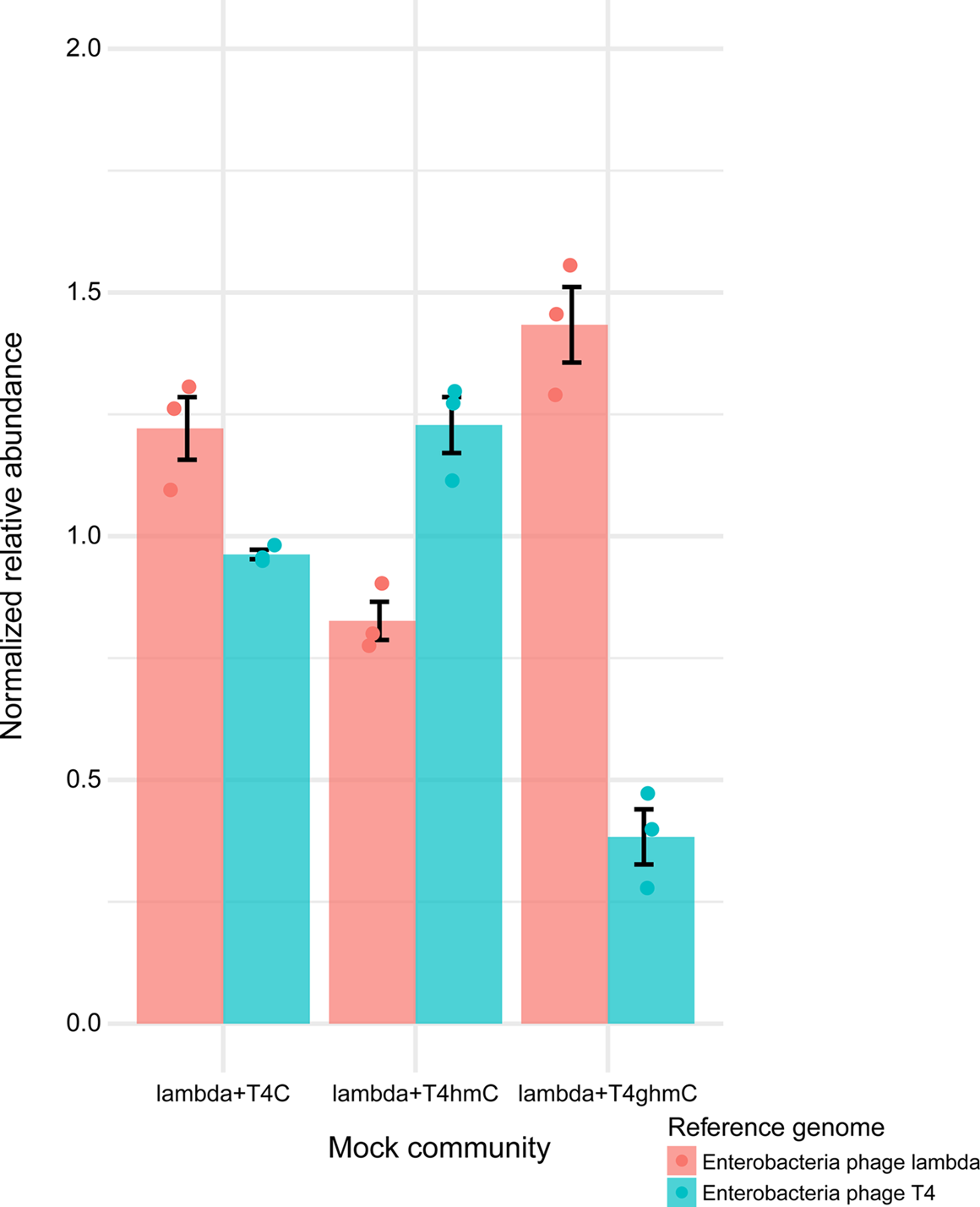
Assessing the potential inhibitory effect of gluycosyl-hydroxymethylcytosine and hydroxymethylcytosine on recovery of modified T4 viral genomes. T4 DNA with (hmC, ghmC) or without (C) base modifications was mixed with lambda and subjected to Illumina sequencing. The graph shows the relative abundance of T4 and the lambda control by RPKM from sequencing normalized to that of the input genome copy number from qPCR. ghmC: glucosyl-hydroxymethylcytosine. hmC: hydroxymethylcytosine.

## Discussion

Here we describe steps toward optimizing virome purification methods, in part by taking advantage of tests with a mock community comprised of eight diverse viruses (VirMock1). Previously multiple groups have published studies of optimizing methods for virome analysis[8, 18, 28-33, 43-45]. All of these are valuable, but in any biochemical manipulation there are typically multiple alternatives at each step, so that the parameter space is large. Here we work to fill in more of this space with control experiments. In addition, goals in virome studies have been changing over time. Early workers were generally content with recovering large numbers of viruses reproducibly, and were less concerned with capturing every possible virus in a community. As the field has matured, interest has grown in the “unknown unknowns”, and methods optimization has broadened to include difficult to recover viral types that might once have been passed over. At present, multiple large-scale virome programs are initiating, motivating renewed focus on optimizing methods for capturing the widest range of viruses possible.

Some of our main conclusions include the following. Multiple published protocols can enrich virus particles from stool and saliva; the usefulness of enrichment steps for low-biomass samples such as BAL and OP wash is uncertain. Conventional metagenomic DNA sequencing recovered dsDNA viral genomes, and RNA-seq recovered RNA viral genomes, as expected, but the viral-enrichment protocols greatly increased the fraction of reads attributable to contigs annotated as viral. Amplification methods introduce distortions into the data and should be used with caution. Most methods display some degree of over-amplification of M13, with GenomiPhi showing the strongest biases, consistent with several previous studies reporting biases towards small circular ssDNA genomes using MDA [32, 53, 71, 72]. Most viral particles can withstand nuclease treatment, but MS2 was sensitive in some settings; one conjecture is that the structure of the MS2 particle, which includes a portal protein embedded in the icosahedral protein shell, somehow exposes the viral genome to degradation. A longer read Illumina sequencing method allowed recovery of longer viral contigs. DNA modifications found in phage T4 DNA did not disrupt capture in the pipelines used, though the presence of glucosylhydroxymethylcytosine did seem to modestly reduce recovery. More analyses of the effects of additional types of base modifications in DNA and RNA would be valuable.

Based on our assays, we prefer VP1 for analysis of stool samples, and VP4 for analysis of saliva samples. Protocols for each are presented in the Supplementary materials (Supplementary Document S1).

Short-read and long-read sequencing each have their own advantages and limitations. Short read yields a large amount of data at a low cost, but the shorter sequencing length (around 300bp) could present challenges to microbial genome assembly, especially for sequences with repetitive regions or high-GC content[73]. This limitation could be overcome by applying single molecule long-read sequencing, as in Oxford Nanpore or PacBio technologies. However, these can be lower-throughput, less accurate, and more expensive[74]. In addition, many viral genomes are below 5 kb in length, and so might be removed in steps to purify longer molecules. As an initial approach to finding a sequencing platform that balances these factors, we compared the Illumina 1000-cycle kit and 300-cycle kit. The 1000-cycle kit generated more viral contigs of longer lengths and is suited to capturing short viral genomes. It will be useful to assess the utility of the different methods more fully.

Choices of virome enrichment methods are dependent on the sample type and goals of the experiment. For stool and saliva samples, VP1 and VP2 are reasonable as standard methods; VP3 may be useful in more specialized settings. In studies where small circular single-stranded DNA viruses are the target of interest, it may actually be advantageous to use GenomiPhi to bias recovery in their favor, though GenomiPhi is not optimal in other circumstances. Viral particle enrichment is likely useful for analysis of stool and saliva, where rich viral communities are present along with a complex background. For dilute low biomass samples, such as BAL, oropharyngeal swab and nasopharyngeal swab eluate, losses during handling become major factors. Recovery was better for VirMock1 in stool or saliva, indicating a potential carrier effect of the complex sample background. For low biomass samples, direct nucleic acid sequencing without enrichment may be best; optimization tests are ongoing.

Multiple additional methods can be used for virome analysis, each of which has strengths and weaknesses. Hybridization capture is one useful method [75], in which a large library of oligonucleotides are generated complementary to viral sequences, and these are used to capture and enrich viral sequences from a complex mixture prior to sequencing. This can be useful when scanning for known viruses, for example in identifying a virus of human cells responsible for a disease outbreak. This method is not useful for studying viruses that are not complementary to the capture primers, commonly including bacteriophages.

Some of the most extreme variations in viral genomes and related genetic elements were not examined here, and would be a useful topic for future methods development. Some of the giant viruses (Phylum Nucleocytoviricota) have huge particle sizes and genomes [39], and may be lost during handling in many virome enrichment protocols. We included vaccinia as a representative, but other Nucleocytoviricota are much larger and may well be lost selectively during handling. At the other extreme, small satellite RNA viruses (such as hepatitis delta and viroids) are small circular RNAs of only a few hundred bases. Recent work has identified such small RNAs widely in diverse sample types[41]. Here we did not attempt to assess capture of either the giant viruses or tiny satellites; further work to quantify recovery of these elements would be useful.

In summary, where possible, it is useful to enrich virome specimens for particles before sequencing to optimize capture of previously unknown viral lineages. This work presents checks on a variety of steps, taking advantage of samples spiked with a community of known composition. However, there are usually multiple reasonable alternatives at each step; here we review outcomes over a variety of protocol choices, but many more alternative methods could reasonably be considered. It would be useful if multiple groups carrying out such methods optimization studies would publish their work to accelerate further technology refinement.

## Methods

### Materials and reagents used

Key reagents are listed in Table S7.

### Virus propagation

Phage T4, lambda, MS2, and phi6 were propagated in their respective host (Table 1, Table S1), except for T4 C, which was propagated in *E. coli* CR63 first but then had a single plaque picked to infect *E. coli* DH10B [49]. All phage lysates were each made from a single plaque, harvested from liquid culture and treated with chloroform. Plaque assays were performed to determine phage titers. M13 was purchased from AntibodyDesignLaboratories (PH010S) and propagated in *E. coli* C3000 to validate titers. AAV-GFP vectors were prepared as described[76]. Briefly, 293T cells were transfected with AAV *trans* plasmid (pAAV-GFP), AAV *cis* plasmid (pAAV2/8), and helper plasmid (pAdDeltaF6), then incubated overnight before washed and incubated for another 48 hours prior to isolation by centrifugation, nuclease treatment, and chloroform treatment. AAV titers were determined by GFP expression measured Incucyte live imaging analysis and by qPCR. Vaccinia virus (strain Western Reserve, ATCC VR-119) was propagated by infecting BSC-1 cells were with VV with MOI of 1:10 and then incubating at 37°C for 48 hours. Viruses were then harvested by freezing the cell suspension at -80°C, repeating freeze-thaws for 3 times followed by sonication of the cells, and taking the supernatant. Titers of VV were quantified by crystal violet staining. Murine hepatitis virus was provided by Dr. Susan Weiss. Briefly, MHV was propagated in cultured 17Cl-1 cells and quantified for titers by crystal violet staining.

### Plaque assays

Bacterial hosts were cultured overnight. OD600 was measured and then bacterial cultures were mixed with 0.5% top agar and subsequently poured onto LB agar plates. Serial dilution of phages were then plated onto bacteria in separate spots. Plates were incubated at 37°C overnight before counting plaque-forming units. VV was plaqued on BSC-1 cells. Crystal violet staining was used to determine the plaque-forming units (pfu) of VV. MHV was plaqued on L2 cells. Crystal violet staining was used to determine the pfu of MHV.

### qPCR

All qPCR assays for reference viruses in VirMock1 were carried out using the TaqMan Fast Virus 1-Step Multiplex Master Mix (No ROX) with the following conditions: 5min at 50°C, 20s at 95°C and 40 cycles of 3s at 95°C and 30s at 60°C. qPCRs were performed on a QuantStudio 5 Real-Time PCR System. qPCR probes and primers were ordered from IDT as pre-mixed assays (Table S8).

Plasmids based on pUC57 with PCR amplicons inserted were purchased from GenScript and used as qPCR standards for all viruses except AAV (Table S9). The standard for AAV was trans pAAV-GFP plasmid (Addgene plasmid #32395).

### Characterization of modified T4 DNA with by digestion with the restriction endonuclease Alu I

DNA was extracted from 140 μL of stock T4ghmC, T4hmC and T4C using QIAamp RNA viral mini kit generally following manufacturer’s instructions, except for the use of RNase-free water instead of Buffer AVE for elution. DNA extracted from T4ghmC, T4hmC, and T4C was digested with C-specific AluI restriction enzymes at 37°C for 1 hour and then visualized on 1% agarose gel electrophoresis with ethidium bromide (Fig. S4).

### Liquid chromatography-mass spectrometry (LC-MS) analysis to characterize modified T4 DNA

DNA samples were extracted as mentioned above for AluI digestion. Extracted DNA samples were denatured at 95 °C for 5 min. The resulting single-stranded DNA was digested with nuclease P1 (New England Biolabs, M0660S) in the appropriate digestion buffer at 37 °C for 2 h, followed by digestion with calf intestinal alkaline phosphatase (CIP, New England Biolabs, M0525S) at 37 °C overnight. The digested nucleoside solutions were diluted to a final concentration of 2.5 ng/L. Formic acid was added to all samples to a final concentration of 0.1% prior to LC–MS analysis.

Liquid chromatography was performed on an Ultimate 3000 UPLC system (Thermo Fisher Scientific) using Buffer A (1% acetonitrile, 0.1% formic acid in H₂O) and Buffer B (0.1% formic acid in acetonitrile) at a flow rate of 300 nL/min. One microliter of each sample was loaded onto a reverse-phase C_18_ column (Acclaim PepMap 100, Thermo Fisher Scientific, 164534) maintained at 30°C. The separation was achieved using a segmented linear gradient of Buffer B as follows: 1% from 0–2 min, ramping to 40% at 4 min, increasing rapidly to 95% at 5 min, held at 95% from 5–6 min, and returned to 1% at 7 min for re-equilibration. Eluted nucleosides were introduced into the mass spectrometer (Exploris 240, Thermo Fisher Scientific) via nano-electrospray ionization using NanoESI emitters (FOSSILIONTECH, The Sharp Singularity) operated at 2500 V. Parallel reaction monitoring (PRM) was applied targeting the nucleosides of interest. Full MS^1^ scans were acquired over an m/z range of 200–500 at 120,000 resolution with a maximum injection time of 100 ms. Targeted precursor ions (10 ppm tolerance) were isolated within a 1 m/z window, then fragmented by higher-energy collisional dissociation (HCD) at 25% normalized collision energy. MS^2^ scans were recorded at 60,000 resolution with a dynamic exclusion time of 8 s. Raw data were processed using FreeStyle 1.6 (Thermo Fisher Scientific). Target mononucleosides were confirmed by characteristic product ions in the MS^2^ spectra (Table S10). Quantification was performed based on extracted ion chromatogram (XIC) peak areas with a 5 ppm mass accuracy threshold (Table S11). To compare the relative abundance of modified/unmodified cytosine across the samples, normalization was performed using the signal of 2′-deoxythymidine (dT) to correct for differences in sample loading (Table S11 and Fig. S3). For each nucleoside, normalized relative abundance was calculated as:

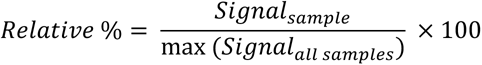

### Assembling the mock viral community

Over the course of the experiments, multiple batches of VirMock1 were assembled. VirMock1 was prepared by mixing T4 ghmC, lambda, MS2, phi6, M13, MHV, AAV vector, and VV. Mixtures were then stored in single-use aliquots at -80°C. See Table S3 for VirMock1 composition of each batch as quantified by qPCR.

### Nuclease titration experiments on VirMock1

1X nuclease included 10X DNase buffer (Roche, 4716728001), 2 μL of Roche DNase I (Roche, 4716728001) and 1 μL of RNase (Roche, 11119915001). For every 100 μL of post-virome prep sample, nuclease digestion was carried out at 75°C for 30 minutes, followed by adding 1 μL of 0.5M EDTA and incubating at 37°C for 10 minutes to inactivate the nucleases.

Nuclease titration experiment using mock community only: Mock community was treated with 0.1X nuclease (10-fold dilution of the DNase I and RNase used in 1X nuclease), 1X nuclease, 10X nuclease (10-fold concentration of the DNase I and RNase used in 1X nuclease), or no nuclease.

Nuclease titration experiment using the mock community spiked into stool: The mock community was spiked into 200mg of solubilized stool, passed through VP1, then treated with 0.1X, 1X, 10X, or no nuclease.

### Nuclease titration on free nucleic acid

Mouse b2m DNA (accession: NM_009735.3) and Rat total RNA (accession: NM_017015.3) were spiked into VirMock1 in a total volume of 150 μL and passed through VP4 with different titrations of nuclease (none, 0.01X, 0.1X, and 1X nuclease). qPCR assays were performed to quantify the amount of mouse b2m DNA and rat total RNA after treatment (Table S4). IDT qPCR assay (ID: Rn. PT39a.22214822.g) was used for quantifying Rat total RNA. See Table S8 and S9 for the qPCR assay used for mouse b2m DNA.

### Base modification experiment

Lambda was mixed with different T4 strains, including T4 ghmC, T4 hmC, and T4 C, in proportions provided in Table S6, and then extracted using the QIAamp Viral RNA kit. The extracted nucleic acid was prepared for sequencing using Nextera XT library prep and sequenced using Nextseq1000 (300-cycles).

### Saliva amplification experiment

3.5 mL of pooled saliva was treated with 1 M DTT for a final concentration of 25mM DTT, incubated at room temperature for 2 min, then centrifuged at 2500g for 1 min at 4°C. The supernatant was passed through a 40um strainer (pluriStrainer Mini 40uM, 43-10040-40). The VirMock1 community was then spiked into 394 μL aliquots of saliva and resuspended in SM buffer for a final volume of 600 μL. The sample was then centrifuged at 2,500g for 10min at 4°C. The supernatant was filtered through a 0.8um pore-size PES filter (Sartorius, VK01P042) by centrifuging at 2000g for 6 mins. The filtrate was transferred into Amicon Ultra-4 Centrifugal Filters (Thermo Fisher Scientific, UFC810096) and spun at 4,000g in 10mins intervals until there was ∼100uL remaining in the Amicon. The concentrate was then treated with 1X nuclease as described above. A total of 140μl VLP preparation was used for viral nucleic acid extraction immediately after nuclease treatment using QIAamp Viral RNA kit. Extracted nucleic acid was stored at -80°C until use.

Total nucleic acid was either reverse transcribed and amplified using the Complete Whole Transcriptome Amplification kit (MilliporeSigma, WTA2) following a protocol from Conceicao-Neto et al.[45] (PMID: 26559140), or reverse transcribed using SuperScriptIII and then amplified with GenomiPhiV3 (Cytiva 25-6601-24) or MALBAC (Yikon Genomics, KT110700110) or ResolveDNA Whole Genome Single-Cell Core Kit (BioSkryb Genomics) following manufacturers’ instructions. The PCR products were purified using QIAquick PCR purification kit and stored at -20°C until use. Samples were sequenced using Illumina Nextera XT library prep followed by sequencing on NextSeq2000, except for samples undergoing PTA, which used the library prep from the manufacturer following manufacturer’s instructions and subsequently sequenced on a Miniseq.

### Collection and pooling of human samples

All human subjects research was conducted under protocols approved by the University of Pennsylvania Institutional Review Board (IRB). The samples were collected under multiple protocols, including: #849982, #842613, #810851, #851540, #828422, #834299, and #814859. All samples were collected after informed consent. OP wash involved subjects swishing and gargling 10 ml of 1% saline. Saliva was collected (approximately 5 ml) by subjects spitting into a sterile container. Lower respiratory tract samples, specifically Bronchoalveolar Lavage (BAL), were obtained via bronchoscopy (Protocol #810851). To minimize contamination, upper respiratory tract samples were collected prior to anesthesia, and for bronchoscopy, the first lavage return was discarded.

### Stool solubilization

Frozen stool was thawed processed as described by Dillon et al. 2021[77]. Briefly, stool was diluted 1:1 with dPBS in a 50 ml conical tube with twelve 1 mm glass beads (Thermo Fisher Scientific, AC465941000) before being mixed vigorously for at least 30 seconds or until homogenized. Materials were strained using a 500 um cell strainer (VWR International, 43-50500-03) before being pooled in a sterile container pooling more than 5 donor fecal samples together.

### Methods for purification of virus-like particles from stool

VP1: this protocol follows the protocol published by Liang et al. [44] . Briefly, 200mg of solubilized stool was either spiked or not spiked with mock community, resuspended in 10 mL of SM buffer (50 mM Tris-HCl pH 7.5, 100 mM NaCl, 8 mM MgSO_4_), spun down and filtered through a 0.22-μm-pore-size vacuum filter (Thermo Fisher Scientific, SCGP00525). The filtrate was concentrated using a 100-kDa-molecular-mass Amicon Ultra-15 Centrifugal Filter (Thermo Fisher Scientific, UFC910096), resuspended in 10 ml SM buffer and concentrated for the second time to a final volume of around 250μl. The concentrate was topped up to 400 μL using SM buffer and then treated with 1X nuclease as described above. A total of 500μl VLP preparation was used for viral nucleic acid extraction immediately after nuclease treatment using QIAamp Viral RNA kit (per manufacturer, extracts total nucleic acids). Extracted nucleic acid was stored at -80°C until use.

VP2: This protocol was adapted from Conceicao-Neto et al. [45]. Briefly, 50mg of solubilized stool was either spiked or not spiked with mock community, resuspended in 500 μL of SM buffer, and centrifuged at 17,000g for 3 mins. The supernatant was passed through a 40um pore-size strainer (pluriStrainer Mini 40uM, 43-10040-40) and then a 0.8um pore-size PES filter (Sartorius, VK01P042). The filtrate was topped up to 500 μL using SM buffer and then treated with 1X nuclease (as described above). A total of 600μl VLP preparation was used for viral nucleic acid extraction immediately after nuclease treatment using QIAamp Viral RNA kit. Extracted nucleic acid was stored at -80°C until use.

VP3: this protocol follows Shkoporov et al. [43]. Briefly, 500 mg of solubilized stool was either spiked or not spiked with mock community, and then resuspended in 10mL SM buffer. The resuspended sample was homogenized by vigorous vortexing, centrifuged twice, then filtered twice through a 0.45 um pore PES syringe-mounted membrane filters (Millipore, SLHPR33RS). NaCl and PEG-8000 powders were added to reach a final concentration of 0.5M and 10% w/v respectively. Samples were incubated overnight. On the next day, samples were centrifuged to harvest pellets, which were then resuspended in 400 μL SM buffer and equal volume of chloroform to extract. Emulsions were then centrifuged at 2500g for 5 min to harvest the aqueous phase, which was then mixed with 40 μl of a solution of 10 mM CaCl_2_ and 50 mM MgCl_2_, treated with 8U TURBO DNase (Ambion/ThermoFisher Scientific) and 20 U of RNase I (ThermoFisher Scientific), and incubated at 37°C for 1h followed by 10 min at 70°C. Proteinase K (40 μg) and 20 μl of 10% SDS was then added to the tubes and incubated for 20 min at 56 °C, followed by addition of 100 μl of Phage Lysis Buffer (4.5 M guanidinium isothiocyanate, 44 mM sodium citrate pH 7.0, 0.88% sarkosyl, 0.72% 2-mercaptoethanol) and incubation at 65°C for 10 min. The lysates were extracted twice by vortexing with equal volume of Phenol/Chloroform/Isoamyl Alcohol 25:24:1 (Fisher Scientific) and centrifugation at 8000 g for 5 min at room temperature. The nucleic acid was purified from the aqueous phase using DNeasy Blood & Tissue Kit (Qiagen).

VP4: VLP preparation from saliva, OP wash, and BAL Pooled saliva was pre-treated with 1M DTT for a final concentration of 25mM DTT and incubated at room temperature for 2 min. Pooled OP wash and BAL was directly thawed on ice. 557uL of saliva/OP wash/BAL was either spiked or not spiked with mock community and resuspended in SM buffer to reach a final volume of 600 μL. The sample was then centrifuged at 2,500g for 10min at 4°C. The supernatant was filtered through a 0.8um pore-size PES filter (Sartorius, VK01P042) by centrifuging at 2000g for 6 mins. The filtrate was transferred into Amicon Ultra-4 Centrifugal Filter (Thermo Fisher Scientific, UFC810096) and spun at 4,000g in 10mins intervals until there were ∼100 μL remaining in the Amicon. The concentrate was then treated with 1X nuclease as described above. A total of 140μl VLP preparation was used for viral nucleic acid extraction immediately after nuclease treatment using the QIAamp Viral RNA kit. Extracted nucleic acid was stored at -80°C until use. 0.3ng DNA was used for PTA following the manufacturer’s instructions. PCR product was purified using QIAquick PCR purification kit according to the manufacturer’s instructions and stored at -20°C until use.

### Extraction and reverse transcription of viral nucleic acids

DNA and RNA was extracted from purified VLPs using QIAamp Viral RNA extraction kit (Qiagen) following the manufacturer’s instructions. Reverse transcription was then performed on total nucleic acid using SuperScript III (ThermoFisher Scientific). Downstream sequence analysis was performed on DNA and cDNA together.

### Illumina 300-cycles sequencing

All sequencing was performed using Illumina sequencing machines. The Nextera XT library preparation kit (FC-131-1096) was used for all samples. Libraries were tagged using IDT DNA/RNA UD Indexes and quantified with the Quant-iT PicoGreen dsDNA assay. Libraries were sequenced on the same NextSeq 2000 instrument using P1 300-cycle reagents. All libraries received a 1% PhiX spike-in.

### Illumina 1000-cycles sequencing

The library pool as prepared for 300-cycle sequencing had an average fragment size of 542 bp. A 0.55X bead clean-up was performed on the pool using AMPure XP beads (Beckman-Coulter) to target larger fragments and bring the average fragment size closer to 1000 bp. The pool was sequenced on a 2x500 cycle kit on the Illumina MiSeq i100 (the only compatible instrument at the time of the experiment).

### Whole DNA metagenomic sequencing

DNA was extracted from approximately 200 μL total of starting material using the Qiagen DNeasy PowerSoil Pro kit. Extracted DNA was quantified using the Quant-iT PicoGreen dsDNA assay kit (Thermo Fisher Scientific). Shotgun libraries were generated from 7.5 ng DNA using the Illumina DNA Prep Library Prep kit and IDT for Illumina unique dual indexes at 1:4 scale reaction volume. The Quant-iT PicoGreen dsDNA assay kit was used to assess library success. An equal volume of library was pooled from every sample and sequenced using a 300-cycle Nano kit on the Illumina MiSeq. Libraries were then re-pooled based on the demultiplexing statistics of the MiSeq Nano run. The final library pool was QC’ed on the Qiagen QIAxcel with the QIAxcel DNA High Resolution kit to check fragment size distribution and absence of adaptor fragments. Libraries were sequenced on an Illumina Novaseq X flow cell, producing 2x150 bp paired-end reads. Extraction blanks and nucleic acid-free water were processed along with experimental samples to empirically assess environmental and reagent contamination. A laboratory-generated mock community consisting of DNA from *Vibrio campbellii* and Lambda phage was included as a positive sequencing control.

### RNA-seq Analysis

RNA was extracted from approximately 200 μL total of starting material using the Qiagen RNeasy PowerFecal Pro kit. RNA was quantified using the Qubit™ RNA HS Assay Kit (Thermo Fisher Scientific). RNA quality and RIN were checked with the Agilent TapeStation RNA ScreenTape Analysis kit. Libraries were generated from 25-500 ng RNA using the Illumina Stranded Total RNA Prep, Ligation with Ribo-Zero Plus Microbiome and Illumina RNA UD indexes. Library success was assessed by the Quant-iT PicoGreen dsDNA assay kit. Libraries were QC’ed on the Qiagen QIAxcel with the QIAxcel DNA High Resolution kit to check fragment size distribution and absence of adaptor fragments. Libraries were then diluted to the same molar concentration and pooled in equal amounts. Libraries were sequenced on an Illumina Novaseq X flow cell, producing 2x150 bp paired-end reads. Extraction blanks and nucleic acid-free water were processed along with experimental samples to empirically assess environmental and reagent contamination. A positive sequencing control was used consisting of *E. coli* total RNA (Thermo Fisher Scientific).

### Bioinformatic analysis

Sequences were analyzed using the bioinformatics pipeline Sunbeam v4.0.0, comprised of adapter trimming (Trimmomatic), host read removal, quality filtering (Komplexity), taxonomic assignment (Kraken), contig assembly (Megahit v1.2.9), ORF prediction (Prodigal), and virus identification and annotation (Cenote-Taker2 v2.1.3, Cenote-Taker3 Database 3.1.1). To calculate the relative abundance of each virus in the mock community, all reads were trimmed to remove adaptors and filtered with phred quality ≥Q30 using fastp, mapped using Burrows-Wheeler Aligner (BWA) against the reference genomes of all VirMock1 community members (Table S12), and analyzed for the number of mapped reads per kilobase of genome target per million sample reads. The percentage of viral reads was calculated by dividing the total number of sample reads by the number of reads that were mapped to the viral contigs identified by Sunbeam for all experiments.

All plots were generated using RStudio (version 2025.05.1+513) and processed by Illustrator. Bray-Curtis distance calculations and PCoA analysis were carried out in RStudio using the Vegan package. Data for generating the plots is in Table S13.

## Data availability

Sequences used as reference genomes are listed in Table S12. Sample information and raw sequences are available in the National Center for Biotechnology Information Sequence Read Archive under BioProject ID XXX (Table S14).

Raw LC–MS/MS data for DNA nucleoside composition analysis (enzymatic hydrolysis of genomic DNA into individual nucleosides) were deposited in the MassIVE repository under accession number MSV000099555.

All bioinformatics scripts used are deposited at https://github.com/jduan7/VLP-enrichment-methods-optimization-2025.

## Acknowledgements

We are grateful to members of the Bushman and Collman laboratories for help and suggestions, and to participants who provided samples. Thanks to Laurie Zimmerman for help with art work. We thank Dr. Goulian, Dr. Isaac, and Dr. Weiss for providing phage phi6, vaccinia virus, and murine hepatitis virus, respectively. We thank Illumina for making the 1000 cycle kit available for our work.

## Author Contributions

JD, ADM, RGC, and FDB conceived the study; YZ and AT carried out mass spec analysis, JD, ADM, MH, YH, AF and KL carried out biochemical analysis; SH, AMM and NW carried out metagenomic DNAseq and RNAseq analysis; JD, ADM, and NW carried out bioinformatic analysis; JD, ADM, RGC, and FDB wrote the paper.

## Funding

Funding: This work was supported in part by The PennCHOP Microbiome Program, and NIH grants U54AG089323 (Human Virome Program Penn Virome Characterization Center), P30AI045008 (Penn Center for AIDS Research), U19AI174998, and R01LM014503.

## Conflicts of Interest

The authors declare that they do not have any conflicts of interest.

## Supplementary Material

Table S1. Genotype information of T4 variants used in this study.

Table S2. Summary of features of tested amplification methods.

Table S3. Genome copy number of reference viruses in each batch of VirMock1 as quantified by qPCR.

Table S4. qPCR CT values of free mouse b2m DNA and free rat total RNA under nuclease titration (none, 0.01X, 0.1X, 1X).

Table S5. Comparison of 300-cycle kit vs 1000-cycle kit regarding median lengths of total and viral contigs with and without normalization to toal bases sequenced per sample.

Table S6. Relative abundance of lambda and T4 with different base modifications in lambda-T4 mixtures as quantified by RPKM via sequencing and by genome copy number via qPCR.

Table S7. Key reagents.

Table S8. qPCR probes and primers used in this study.

Table S9. Sequences of plasmid and oligos used as standard controls for qPCR assays.

Table S10. Mass of nucleosides and product ions by LC-MS.

Table S11. Quantification of mononucleosides in T4ghmC, T4hmC and T4C by LC-MS.

Table S12. Accession numbers for reference sequences used in this study.

Table S13. Input data files for R scripts used to generate the figures in this study.

Table S14. Metadata and accession numbers for synthetic samples used in this study.

Supplementary Document S1. Complete workflow of our favored virus enrichment protocols (VP1 and VP4)

## Supplementary Figures

Figure S1. Relative abundance of class of viruses as annotated by Cenote-Taker2 in saliva, stool or BAL that was analyzed by direct extraction of DNA (red) or RNA (blue), or after viral particle enrichment and analysis using VP1, VP2, VP3, or VP4.

Figure S2. Relative abundance of each reference virus after VirMock1 was spiked into saliva, OP wash, or BAL and followed by VP4 and amplification by PTA or remained unamplified.

Figure S3. Validation of DNA modifications in T4 strains by LC-MS. Relative abundance of each modified and unmodified nucleoside in T4 C, T4 hmC, and T4 ghmC determined by LC-MS are shown in the bar graph.

Figure S4. Validation of T4 strains by AluI digestion of T4 DNA. Phage T4 ghmC, T4 hmC and T4 C DNA left untreated (-) or treated (+) with the restriction enzyme AluI, which cleaves unmodified DNA.

## Notes

### Competing Interest Statement

The authors have declared no competing interest.

