## Supplemental Document S1 for "Optimizing methods for virome analysis based on studies of a synthetic viral community"

### Supplementary Document S1. Complete workflow of our favored virus enrichment protocols (VP1 and VP4)

**VP1: used for enrichment of virus-like particles from stool samples**

<u>Isolation of virus-like particles</u>

1. Thaw 200mg of stool and resuspend in 10mL of SM buffer. Vortex to mix well.
2. Centrifuge the sample at 2,500g for 10min at 4C.
3. Take the supernatant and pass through the 0.22um Sterilflip vacuum PES filter into a new 15mL conical tube.
4. Briefly spin down at 4,000g for 30s.
7. Pre-wash Amicon Ultra-15 Centrifugal Filter by adding 15mL of 0.45um-filtered SM buffer.
    a. SM buffer: 50mM Tris-HCl pH7.5, 100mM NaCl, 8mM $MgSO_4$
8. Spin for 5 mins at 4,000g. Decant the flow through.
9. Transfer the filtrate from step 4 into the Amicon.
10. Spin at 4,000g in 10mins intervals until there are ~250uL remaining in Amicon.
11. Resuspend samples in 10 ml SM buffer and concentrate for the second round (4,000g at 10mins intervals) to a final volume of around 250uL.
    a. The volume that samples could be reconcentrated down to may be variable each time and not necessarily uniform for each sample (usually varying in the range between 100uL to 320uL from our observation). To ensure consistent nuclease treatment, add appropriate volume of SM buffer to top up everything to the same volume as the sample that has the highest volume remaining after two rounds of reconcentration. For the subsequent step, we present an example of topping all sample volumes up to 400uL.
11. Make master mix for DNase & RNase digestion as following:

| Reagent | 1 rxn (uL) |
|---|---|
| 10X DNase buffer | 50 |
| Water | 38 |
| RNase | 4 |
| DNase I | 8 |
| Master mix total | 100 |
| Sample | 400 |
| Total | 500 |

15. Incubate at 37C for <u>30 mins</u>.
16. Add 1uL of 0.5M EDTA into each sample.

17. Incubate the samples at 75C for 10 mins.
18. Proceed to nucleic extraction using QIAamp Viral RNA mini kit on the same day. Store at 4C until use.

**VP4: used for enrichment of virus-like particles from saliva samples**

Pre-filtration of saliva

1. Add 1M DTT to saliva at a final concentration of 25mM.
2. Swirl the tube. Incubate at room temperature for 2 mins.
3. Optional: Centrifuge the sample at 2,500g for 1 mins at 4C. Transfer the supernatant through the 40um strainer that is placed on a new 15mL conical tube. Incubate for 5 mins for saliva to pass through.
4. Store on ice until use.

Isolation of virus-like particles

1. Centrifuge culture supernatant at 2,500g for 10min at 4C.
2. Filter the supernatant through 0.8um PES filter.
    a. Add <600uL supernatant into the Vivaclear.
    b. Centrifuge at 2000g for 6 mins.
3. Pre-wash Amicon Ultra-4 Centrifugal Filter by adding 5mL of 0.45um-filtered SM buffer.
    a. SM buffer: 50mM Tris-HCl pH7.5, 100mM NaCl, 8mM $MgSO_4$
4. Spin for 5 mins at 4,000g. Decant the flow through.
5. Transfer the filtrate from step 2 into the Amicon.
6. Spin at 4,000g in 10mins intervals until there are ~100uL remaining in Amicon.
7. Transfer the sample in the Amicon into an Eppendorf tube.
8. Top up the volume with SM buffer to ~100uL.
9. Make master mix for DNase & RNase digestion as following:

| Reagent | 1 rxn (uL) |
|---|---|
| 10X DNase buffer | 14 |
| Water | 23 |
| RNase | 1 |
| DNase I | 2 |
| Master mix total | 40 |
| Sample | 100 |
| Total | 140 |

10. Incubate at 37C for 30 mins.
11. Add 1uL of 0.5M EDTA into each sample.
12. Incubate the samples at 75C for 10 mins.
13. Proceed to nucleic extraction using QIAamp Viral RNA mini kit on the same day. Store at 4C until use.

**Nucleic acid extraction using QIAamp Viral RNA mini kit**

1. Mix 140uL of VLP sample with 560uL of AVL buffer. Vortex.
2. Incubate at rt for 10 min.
3. Briefly spin down.
4. Add 560uL of absolute ethanol to the sample mixture.
5. Mix well by vortexing. Briefly spin down.
6. Add 630uL of the sample mixture into the spin column.
7. Centrifuge at 6,000g for 1 min. Change the collection tube.
8. Repeat step 8-9 until all samples have been loaded.
9. Add 500uL of AW1 buffer to the spin column.
10. Centrifuge at 6,000g for 1 min. Change the collection tube.
11. Add 500uL of AW2 buffer to the spin column.
12. Centrifuge at 20,000g for 3 min. Change the collection tube.
13. Centrifuge at 20,000g for 1 min. Place the spin column in an RNase-free 1.5mL tube.
14. Add 60uL of AVE (elution buffer) to the spin column. Incubate for 1 min.
15. Centrifuge at 6,000g for 1 min.
16. Store viral RNA and DNA at -80C.

**Reverse transcription using the SuperScript III Kit**

1. Clean the PCR hood and all equipment with RNase-OFF. Repeat cleaning with 70% EtOH.
2. Use RNase-free microcentrifuge tube to prepare the RT-PCR.
3. Make the master mix 1 in a 1.5mL tube.

| | uL per rxn |
|---|---|
| 50uM random hexamers (50 ng/uL) | 1.5 |
| 10mM dNTPs mix | 1.5 |
| DDW | 1.5 |
| Total | 4.5 |

4. Aliquot 4.5uL of master mix 1 to PCR strip tubes. Add 15uL of the extracted nucleic acid (DNA and RNA).
5. Heat mixture to 65C for 5 mins.
6. After the incubation, briefly centrifuge the tubes and place on ice for > 1 min.
7. Make the master mix 2 in a 1.5mL tube.

| | uL per rxn |
|---|---|
| 5X First-strand buffer | 6 |
| 0.1M DTT | 1.5 |

| | |
|---|---|
| RNaseOUTTM Recombinant RNase Inhibitor | 1.5 |
| SuperScripteTM III RT (200U/ul) | 1.5 |
| Total | 10.5 |

8. Add 10.5 uL of master mix 2 to each sample in the PCR strip tubes.
    a. Mix by pipetting gently up and down 10 times and spin down.
9. Incubate at 42C for 50 mins, 70C for 10 mins, then hold at 4C.
