## Supplementary figures and images for "Optimizing methods for virome analysis based on studies of a synthetic viral community"

### Supplemental Figures 1-4

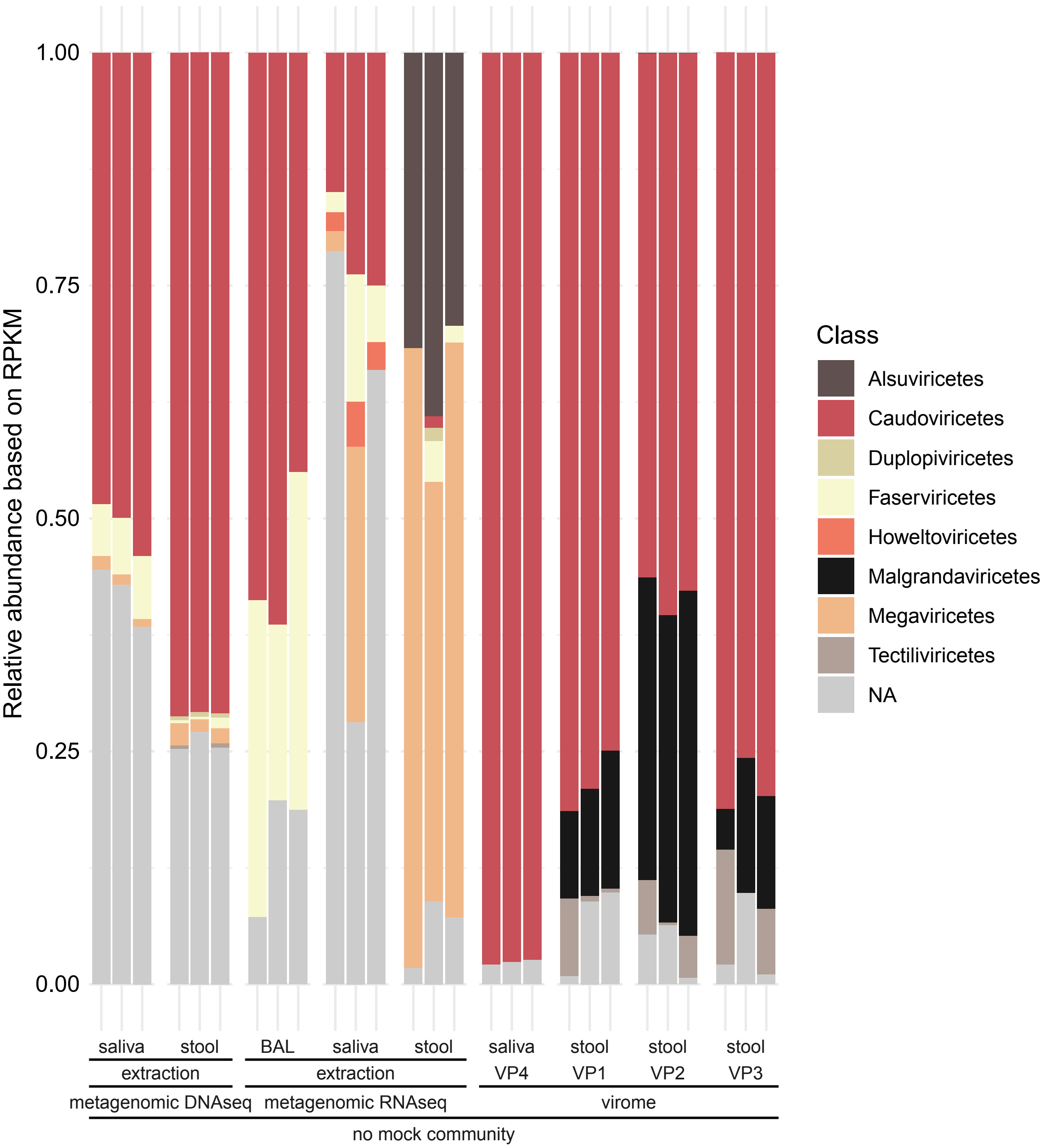

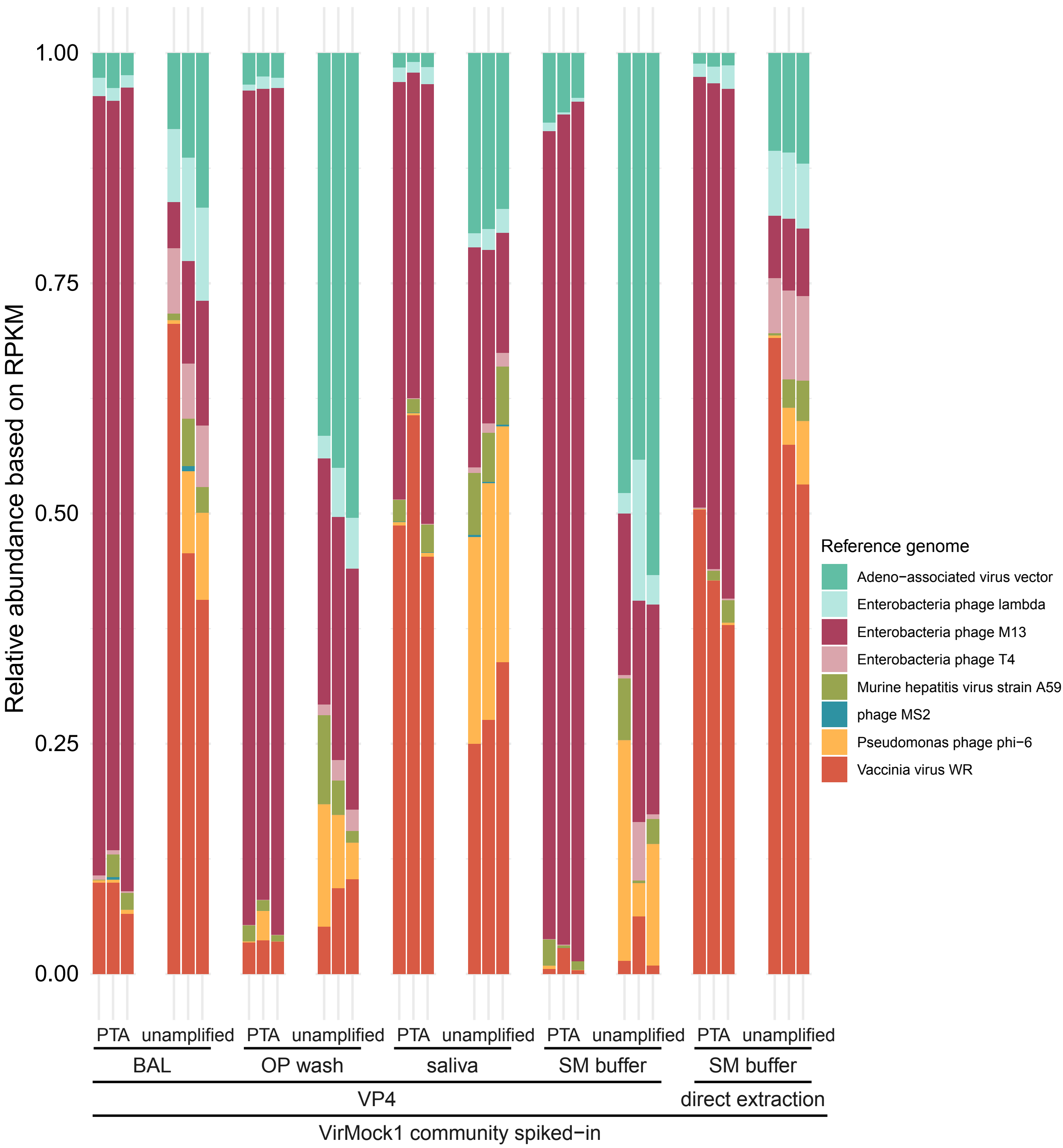

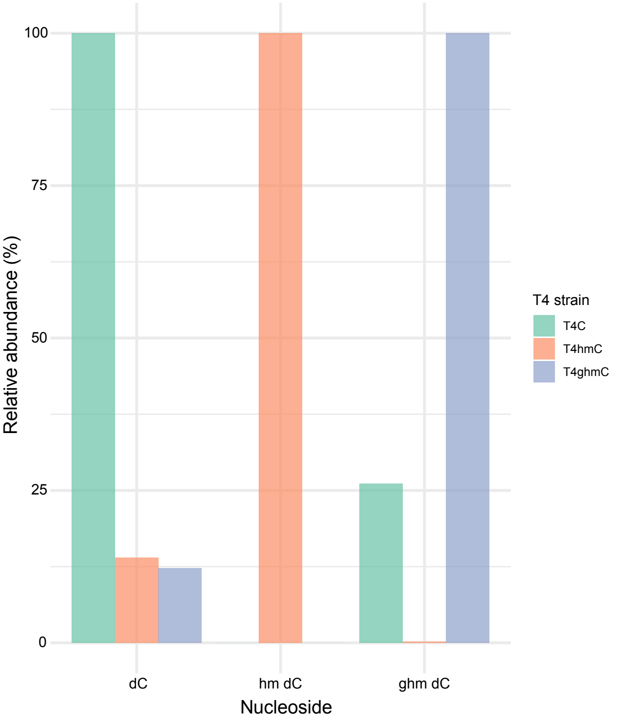

#### Figure S4

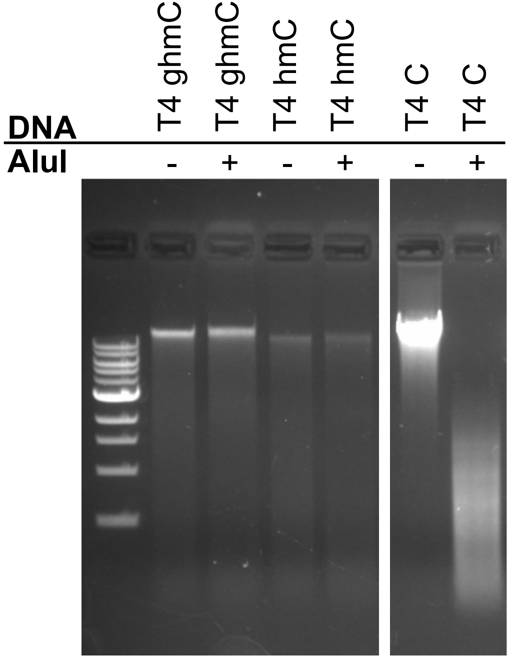
